# Training level reveals a dynamic dialogue between stress and memory systems in birds

**DOI:** 10.1101/2021.01.05.425468

**Authors:** Flore Lormant, Vitor Hugo Bessa Ferreira, Julie Lemarchand, Fabien Cornilleau, Paul Constantin, Céline Parias, Aline Bertin, Léa Lansade, Christine Leterrier, Frédéric Lévy, Ludovic Calandreau

**Author notes:** Correspondence should be addressed to: Ludovic Calandreau, INRAE, UMR 85 Physiologie de la Reproduction et des Comportements, 37380 Nouzilly, France. Shared co-first authorship (Flore Lormant, Vitor Hugo Bessa Ferreira).

## Abstract

It is now well-accepted that memory is a dynamic process, and that stress and training level may influence which memory system an individual engages when solving a task. In this work, we investigated whether and how chronic stress impacts spatial and cue-based memories according to training level. To that aim, control and chronically stressed Japanese quail were trained in a task that could be solved using spatial and cue-based memory and tested for their memory performances after 5 and 15 training days (initial training and overtraining, respectively) and following an emotional challenge (exposure to an open field). While chronic stress negatively impacted spatial memory in chronically stressed birds after initial training, this impact was lowered after overtraining compared to control quail. Interestingly, the emotional challenge reinstated the differences in performance between the two groups, revealing that chronic stress/overtraining did not eliminate spatial memory. Differences caused by previous stressors can re-emerge depending on the more immediate psychological state of the individual. Contrary to spatial memory, cue-based memory was not impaired in any test occasion, confirming that this form of memory is resistant to chronic stress. Altogether these findings reveal a dynamic dialogue between stress, training, and memory systems in birds.

## 1. Introduction

It has long been recognized that chronic stress strongly modulates learning and memory performances [1–3]. However, the relationship between chronic stress and memory are still controversial: some studies reported an improving effect of chronic stress on memory performances [4–6], whereas others reported a negative effect [6–8]. Overall, chronic stress generally provokes memory impairment for complex tasks and improves simple tasks [3]. These discrepancies support the fact that memory is not a single unit. Different parts of the brain are responsible for different memory systems that process different types of information and are differently affected by stress [9].

Two memory systems have had a considerable amount of scientific interest in the last decades as they present strong parallels between humans and animals. The first one is spatial memory, a form of declarative memory based on the hippocampus and used mainly to establish a relationship between different cues from the environment, which allow the creation of a cognitive mental map [10,11]. As the cues from the environment are treated in relation to each other, the spatial memory is considered to be more complex than the cue-based memory, a non-declarative form of memory that results from a simple association between a salient cue and the target and is based on the striatum [9,12,13]. Studies have shown that chronic stress has a strong and negative impact on declarative memory, such as spatial memory [1,3,6,7,14,15], whereas it spares or even improves forms of non-declarative memory [5,6,15,16].

Beyond chronic stress, the training level can also influence the use of multiple memory systems [9,17–20]. Indeed, rats trained to learn the location of a reward in a maze preferentially use their spatial memory after a few days of training and shift to a cue-based dominant response with overtraining [21]. Training and stress levels may also interact: the initial differences in memory performances due to stress may disappear with overtraining [3]. Other studies suggest training itself may be a source of stress and reduce hippocampal neurogenesis [22,23], which could explain the shift between memory systems. While much is known on chronic stress effects on declarative spatial or non-declarative cue-based memory performances, the influence of the level of training and its interaction with chronic stress influence remains poorly understood.

Using the Japanese quail (*Coturnix coturnix*) as the animal model, in this work, we investigated whether and how chronic stress could differentially impact spatial and cue-based memories according to the level of training. Control and chronically stressed Japanese quail were trained in a dual spatial/cued task, which could be solved either using the spatial or cue-based memories and tested for their performances on each type of memory after five and fifteen training days. Additionally, as an animal’s immediate psychological state may interact with chronic stress and impact the relative use of different memory systems and memory performances [24,25], a series of tests were conducted after an emotional challenge that consisted of exposing quail to an unfamiliar environment (open field). Finally, birds were submitted to a test to assess whether they learn to find the target cup by a simple association between the target cup’s colour and the reward or by detecting the difference of colours between the target cup and the remaining cups. Based on the literature, we expected chronically stressed quail to show lower performance in the spatial memory test than control ones after initial training (five days of training). These differences were expected to lower or disappear after overtraining (fifteen days of training) and reinstated after the emotional challenge. Concerning the cue-based memory, we expected no differences between chronically stressed and control quail. Since this is a simple form of memory, we expected the emotional challenge would not disrupt it.

## 2. Methods

### 2.1. Animals

All Japanese quail were bred and maintained at the Pôle d’Expérimentation Avicole de Tours (UE PEAT, INRAE, 2018. Experimental Poultry Facility, DOI: 10.15454/1.5572326250887292E12) where the experiment took place. On the day of hatching, chicks were transferred to communal floor pens. On the 21^st^ day after hatching, chicks were sexed by feather dimorphism, and males were reared in a single home cage (41×51×25 cm) in a battery under a 12:12h light-dark schedule (light on 08h). A piece of artificial turf was placed on the cage floor to allow animals to exhibit dust bathing behaviour. The ambient temperature was maintained at approximately 20±2°C. Unless otherwise specified, food and water were provided *ad libitum*.

Animal care and experimental treatments complied with the French Ministry of Agriculture guidelines for animal experimentation and European regulations on animal experimentation (86/609/EEC). They were performed following the local animal regulation (authorized C37-175-1) of the French Ministry of Agriculture under the EEC directive and under ethics committee approval (Val de Loire, agreement N° 1789 and 1848).

### 2.2. Chronic stress procedure

At 21th day of age, when quail were transferred in individual cages, they were divided into two groups, a control and a chronic stress group. Quail from the chronic stress group were submitted to chronic unpredictable stress (CUS) for 21 days consisting of five to six negative stimulations or stressors per day. To improve the unpredictability of the stressor delivery and decrease habituation to the stress procedure, each stressor occurred at unpredictable times each day during both night and day. The CUS procedure has been fully described and was shown to induce a state of chronic stress in quail [15,26– 30]. Briefly, some stressors were delivered directly in the home cage of the quail: confinement of the bird in the corner of its home cage for 30min, food frustration by placing transparent devices on the feeder just before light on for 30min, soft home cage shaking (2×15 min interspaced by 15 min), unexpected sounds (100 dB) composed of different sounds having or not biological signification for quail. Disturbances from outside the home cage were also delivered, such as a rapid passage of a plastic stick on the rods of the home cage twice a day for 2 min, fast sprayings of water or air on feathers (2 sprays interspaced by 2 min), waving of a plastic flag in front of the home cage for 2 min. Other stressors were also delivered out of the home cage when birds were transferred individually into a new environment for 30 min. Birds from the same line were also placed together in a transport cage for 30 min or placed on a cart and rolled about in the facility.

The control group was left undisturbed except for routine husbandry procedures similar to those provided to birds exposed to CUS. The experimenter regularly visited the control quail room to spend the same amount of time with control and chronically stressed birds. Systematically, when stressors prevented animals from having access to food, an opaque device was placed on the feeder of control birds to prevent them from eating food for an equivalent time.

### 2.3. Learning and memory task

At the end of the CUS procedure, 17 control and 20 chronically stressed birds were submitted to a training procedure followed by a series of memory tests (Figure 1). Briefly, after preliminary phases dedicated to familiarized birds to the food reward (mealworms), the cups and the arena, control and chronically stressed birds were submitted to a dual spatial/cued task during five days that consisted of learning to find food (mealworm) hidden in a single cup (target cup) in an arena that contained eight opaque cups. On each trial, birds were introduced in the arena by a different starting point among three possibilities. The location of the target cup was constant among trials and days of training, allowing animals to solve the task using spatial memory. Moreover, the target cup was black, whereas other non-target cups were white. Thus, birds could also solve this task using their cue-based memory, based on the colour of the cup.

**Figure 1:**
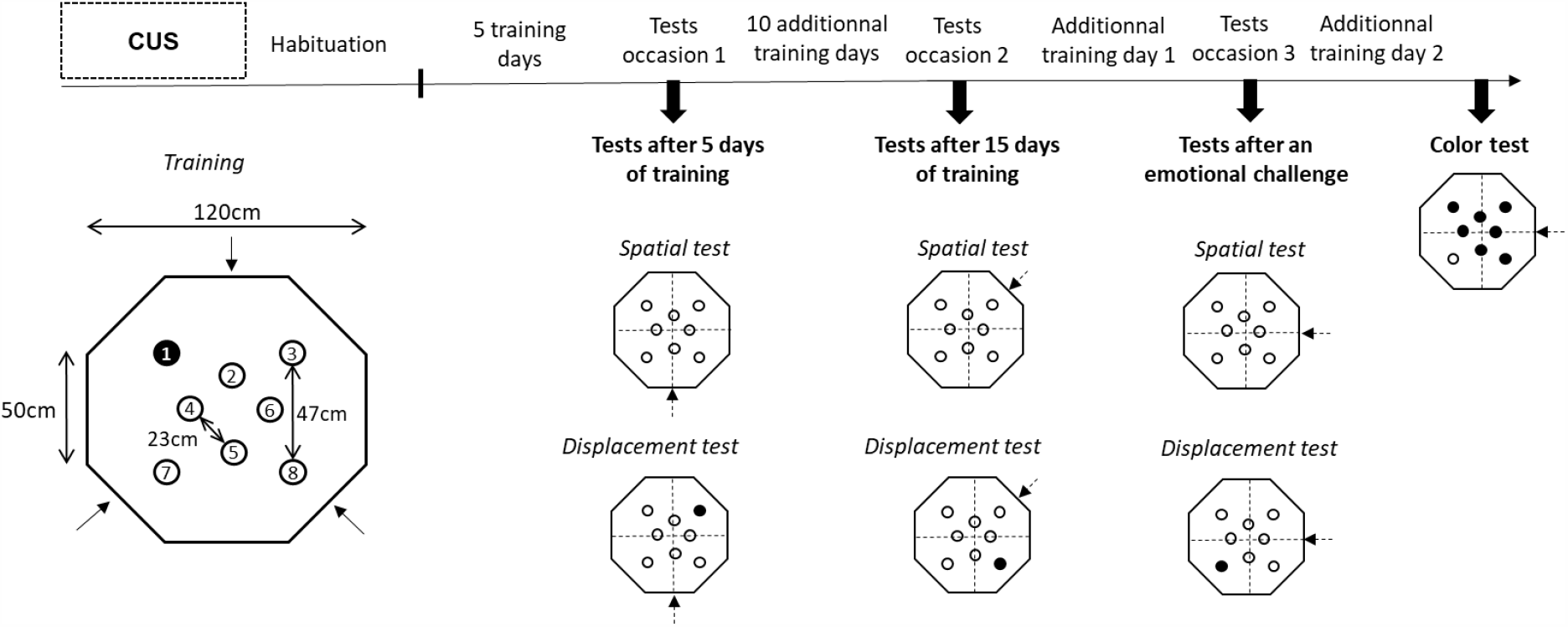
Schedule of the experiment and schematic representation of the apparatus used for training and testing. At three-weeks-old, quail were submitted to chronic unpredictable stress (CUS) for three weeks. At the end of the CUS procedure, stressed and control birds were trained in a dual spatial/cued learning task and submitted to spatial and displacement memory tests after 5 and 15 days of training. After an additional training day, quail were submitted to an emotional challenge (exposure to an open field) followed by a new series of tests. After the last training day, birds were submitted to the colour test. Quail could enter the arena from three different starting points (dark arrows), three starting points during training, and one during testing. The dotted lines represent the division of apparatus into four different quadrants.

The day after the fifth day of training, birds were submitted to a spatial test followed by a displacement test. During the spatial test, all cups were white, and the task could only be solved by remembering the spatial location of the target cup during the training phase. During the displacement test, performed 30 min later, the black target cup was displaced to a new position (different from the position used during training). Thus, both spatial and cue-based memories could provide a solution to the test. Quail could go back either to the spatial location of the target cup during the training phase (spatial memory) or go directly to the displaced black cup (cue-based memory) [15,31,32].

After this first test occasion, birds were trained for ten additional days in the dual spatial/cued task before being submitted to the second test occasion of spatial and displacement tests. Following the second test occasion, birds were trained for an additional day (recall training) in the dual spatial/cued task to avoid that lack of food during the second test occasion reduced bird motivation in subsequent tests. After this first recall training day, all birds were challenged by exposure to an open field test and were submitted to the third occasion of spatial and displacement tests.

On the following day, quail were submitted to a second additional training day (recall training) to keep birds motivated. The day after, birds were submitted to a new test, called “colour test”, to assess whether birds found the target cup by a simple association between its colour (black) and the reward or by learning a more general rule that the target cup was of a different colour compared to the remaining cups.

During all the memory tests, behaviours of quail were recorded by a camera placed above the apparatus and computerized by a tracking video system (Ethovision XT; Noldus IT, The Netherlands).

### 2.4. The dual spatial/cued task

The following procedures and the arena used for the task were previously used and validated by our group in chickens and quails [13,15,32,33].

#### 2.4.1. Mealworms familiarization

One week before the beginning of training, a brown ceramic cup (6×7cm) was placed in each quail’s home cage. For seven days, three times per day (spaced about 3h), 3 or 4 mealworms were deposited in the cup to familiarize birds with the cups and mealworms.

#### 2.4.2. Habituation

The day following the end of the familiarization phase, birds were submitted to the habituation phase. One hour before each habituation session, food was removed from the home cage of the animals. During five days, once a day, each quail was introduced into the centre of a beige octagonal arena (120cm long; 50cm high) surrounded by a blue curtain (1.90m high) and lighted by a bulb at the ceiling (18 Lux). Four black spatial cues were placed on the walls of the arena and four others on the curtain. Eight ceramic cups similar to those used for the familiarization were placed in the arena and contained mealworms. Four cups were covered with white paper, whereas black paper covered another four cups. Each bird was allowed to explore the arena and the cups in each habituation session until it found and ate all mealworms or after a maximum time of 600 s. Between each quail and day of habituation, the position of black and white cups was randomly moved. On each day, the number of cups visited was recorded to estimate the level of exploration and habituation of the birds (Figure 1).

#### 2.4.3. Training

On each day of training, birds were submitted to 2 training trials per day with an inter-trial interval of 1 hour. On each trial, birds were introduced in the arena by a different starting point among three possibilities (Figure 1). Only one cup was rewarded and contained three mealworms (target cup). The target cup was always black and at a fixed position on each trial, whereas the seven other cups were non-rewarded and white. The trial ended when the bird reached the target cup and ate the mealworms, or after a maximum time of 300 s. If the bird did not find the target cup, it was gently guided to it and allowed to eat the mealworms before being removed from the arena. Between each trial, the bird returned to its home cage. The latency to find and eat the food in the rewarded black cup was recorded on each trial.

#### 2.4.4. The spatial test

This test was used to assess the impact of chronic stress on spatial memory. Each bird was allowed to freely explore the arena and the 8 cups for 1 minute. All cups were empty and, more importantly, of similar colour (white). The number of cups visited (number of errors) before reaching the target cup was scored. Moreover, the arena was virtually divided into four equal quadrants, and the time spent in each quadrant (spatial quadrant and three other quadrants) during the test was measured using the tracking video system (Figure 1). The time spent on the three other quadrants was averaged.

The spatial test was conducted after 5 and 15 days of training (initial training and overtraining, henceforth) and after an emotional challenge by exposing birds to an open field (emotional challenge, henceforth).

#### 2.4.5. The displacement test

The displacement test was used to assess the impact of chronic stress on the use of spatial and cue-based memory systems. It was systematically conducted 30 min after the spatial test. During this test, similarly to the training phase, the arena was equipped with one black cup and seven white cups, but all cups were empty of food. The black cup was displaced to a new position (Figure 1). The new position was changed for each test conducted after initial training, overtraining, and emotional challenge. All birds were allowed to explore the arena and the cups for 1 min freely. The number of errors before reaching the spatial and displaced cup was scored. Similar to the spatial test, the arena was virtually divided into four equal quadrants (spatial quadrant, displaced cup quadrant, and two other quadrants), and the time spent in each quadrant during the test was measured using the tracking video system. The time spent on the two other quadrants was averaged.

#### 2.4.6. Tests after an emotional challenge

The third occasion of both tests (spatial and displacement tests) was conducted after an emotional challenge. This emotional challenge consisted of exposing the birds to an open field test, which is known to provoke anxiety-like quail behaviors [15,34,35]. Each quail was individually placed in the centre of a square arena (80 × 80 × 80 cm) of white wood with a beige linoleum floor. The arena was placed into a new experimental room, surrounded by a green curtain and strongly lighted (50 lux). Each quail was allowed to explore the arena for 5 min before being withdrawn from the arena. Thirty minutes after the open field test, quail were submitted to a spatial test followed by a displacement test.

#### 2.4.7. The colour test

This test aimed to assess whether birds learn to find the target cup by a simple association between its colour (black) and the reward or by detecting the difference of colours between the target cup and the remaining cups (Figure 1). Each bird was allowed to freely explore the arena and the eight cups for 1 minute. All cups were empty and black, except for one white cup and not located in the spatial position used for training. The number of visits before reaching the white cup was scored. Moreover, the number of visits before reaching the black cup located in the spatial location used during training was scored and compared to those to reach the white cup and other black cups. As for previous tests, the time spent in the four equal quadrants (white cup quadrant, spatial black cup quadrant, and two other black cup quadrants) of the arena was measured. This test was performed after a second recall training day conducted the day after the third test occasion.

### 2.5. Statistical analysis

Data from habituation (number of cups visited) and from training (latency to reach the target cup) were analysed by parametric analyses of variance (ANOVA repeated measures) with stress group (control and chronically stressed quail) as between-subject factors and days (mean values for both training trials within days) as within-subject factors.

For spatial, displacement, and colour tests, the number of errors (or visits) before reaching the target cup(s) and the time spent in the different quadrant of the arena were analysed by ANOVA with groups (control and stressed) as between-subject factors and test occasions (test after initial training, test after overtraining and test after the emotional challenge) as within-subject factors. Additional within-subject factors (e.g., quadrants or cups considered) were also included, dependent on the variables under consideration. When main effects or interactions were significant, analyses were followed by multiple comparisons corrected by Tukey HSD.

When quail did not find a target cup and visited less than 4 cups, they were considered non-explorer. For these birds, a number of 8 errors and an equivalent time of 15 sec in each quadrant were attributed. All statistical analyses were performed using IBM SPSS 21 and R version 3.6.1. Statistical significance was set at *p* < 0.05.

## 3. RESULTS

### 3.1. Effect of chronic stress on habituation and training in the dual spatial/cued task

During habituation, both groups visited more cups and ate significantly more mealworms over days without any differences between control and chronically stress quail (effect of days: F_4,140_=18.15, p<0.001; effect of stress: F_1,35_=0.13, p=0.72; interaction days x stress: F_4,140_=1.11, p=0.35; Figure 2a).

**Figure 2:**
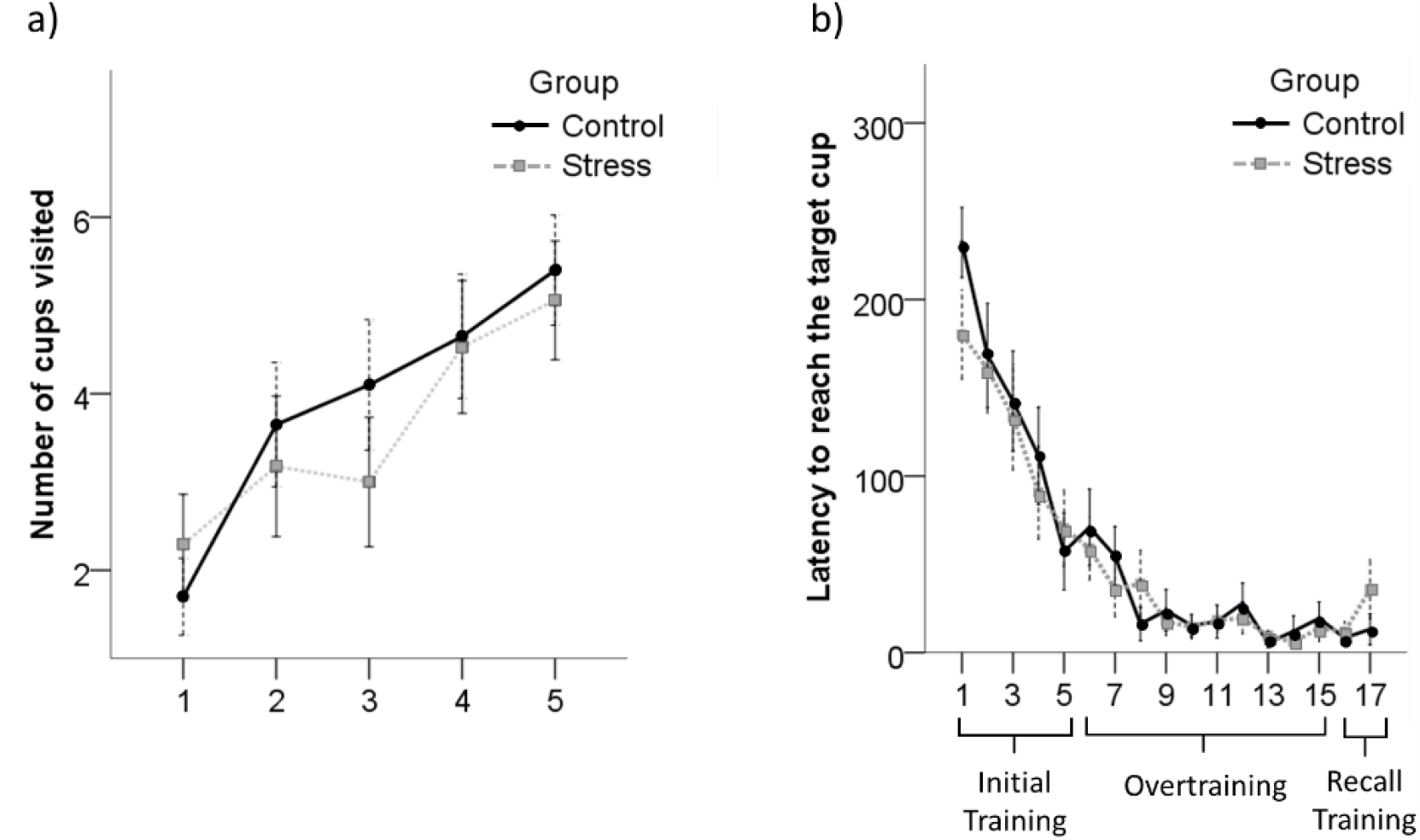
Effects of chronic stress on habituation and training performances in the dual spatial/cued task. **a)** Number of visited cups over days of habituation in control and chronically stressed quail. During habituation, all eight cups were rewarded with mealworms. **b)** Latency to reach the location of the target cup over days of training in control and chronically stressed quail. Mean ± SEM are given.

During the initial training period, both groups similarly learned the location of the target cup since the latency to find the target cup significantly decreased over days (effect of days: F_4,140_=19.60, p<0.001; effect of stress: F_1,35_=0.28, p=0.60; interaction days x stress: F_4,140_=0.93, p=0.45; Figure 2). Similarly, during the overtraining period, the latency to reach the target cup significantly decreased without any effect of chronic stress (effect of days: F_9,315_=5.89, p<0.001; stress effect: F_1,35_=0.13, p=0.73; interaction days x stress: F_9,315_=0.58, p=0.81; Figure 2b).

The latency between the last day of overtraining (the fifteenth day of training) and the recall period between tests (last two additional days) was not different between days, nor between control and chronically stressed quail (effect of days: F_2,70_=1.12, p=0.33; effect of stress: F_1,35_=0.65, p=0.42; interaction days x stress: F_2,70_=1.10, p=0.33), indicating that animals reached a learning plateau.

### 3.2. Effect of chronic stress during the spatial test

The number of errors made before reaching the spatial cup was significantly increased in chronically stressed quail (effect of stress: F_1,35_=5.62, p=0.023, Figure 3a), independently of the test occasion and the interaction with stress (effect of test occasion: F_2,70_=1.45, p=0.24; interaction test occasion x stress: F_2,70_=2.72, p=0.072).

**Figure 3.**
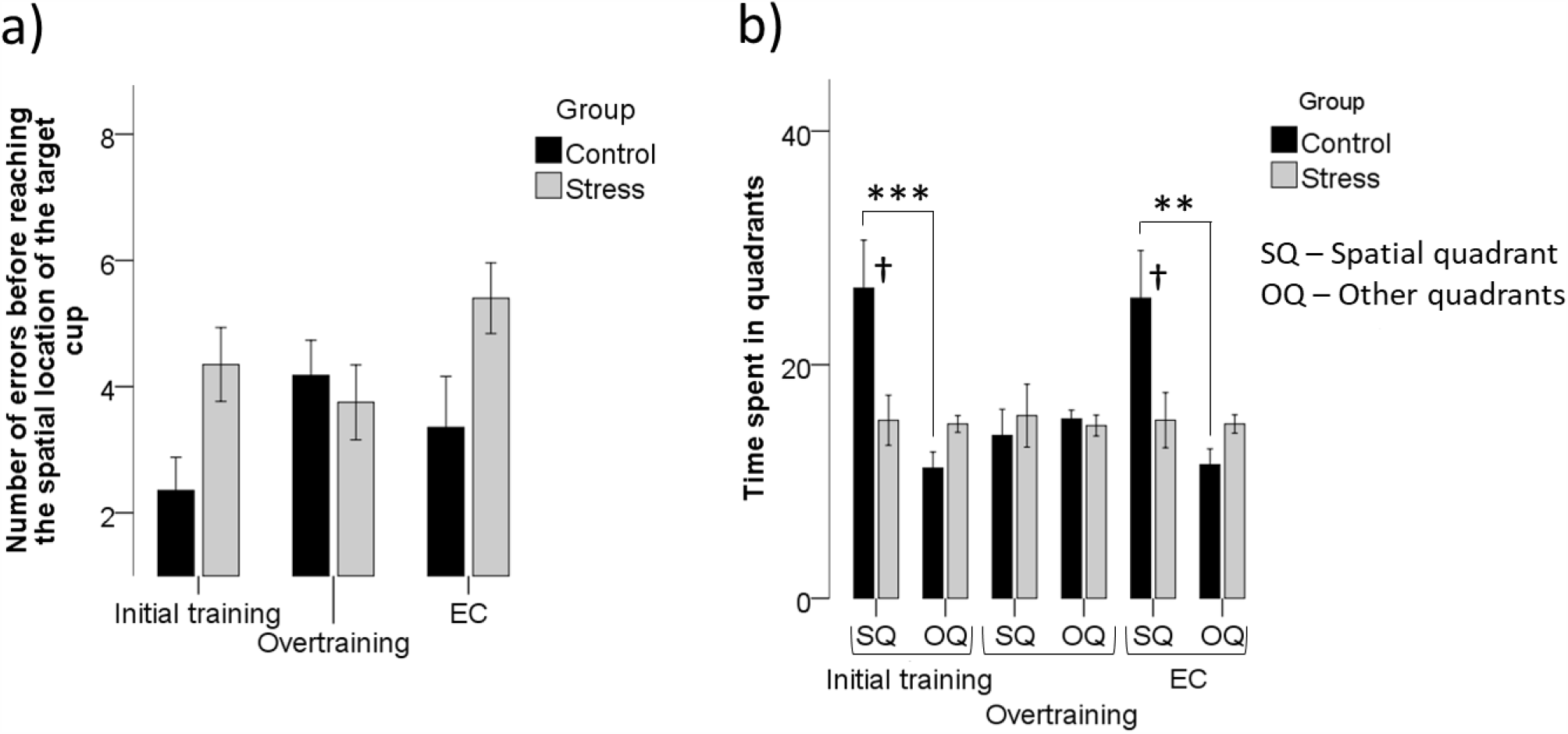
Effects of chronic stress in spatial memory tests performed after initial training (5 days), overtraining (15 days), and after an emotional challenge (EC, exposure to an open field). **a)** Number of errors before reaching the spatial location of the target cup, and **b)** Time (in seconds) spent in different quadrants of the arena (spatial and other quadrants). † p≤0.05, significant difference between control and chronically stressed quail. ** p≤0.01; *** p≤0.001, significant difference between the spatial quadrant and other quadrants. Mean ± SEM are given.

The time spent in the spatial quadrant compared to other quadrants was affected by the quadrant considered, stress, and the interaction quadrant, test occasion and stress (effect of quadrant: F_1,35_=7.58, p=0.009; effect of stress: F_1,35_=6.13, p=0.018; interaction quadrant x test occasion x stress: F_2,70_=3.36, p=0.040). Post-hoc analyses revealed that, during the tests after the initial training and the emotional challenge, control birds spent significantly more time in the spatial quadrant compared to other quadrants (p<0.001 and p=0.001, for the tests after the initial training and the emotional challenge, respectively). In contrast, chronically stressed birds spent an equivalent time in the spatial quadrant compared to other quadrants (p=1 and p=1 for the tests after the initial training and the emotional challenge, respectively, Figure 3b). No differences were found for the test after overtraining: both control and chronically stressed quail spent an equivalent time in the spatial quadrant compared with other quadrants (p=1 and p=1, for control and chronically stressed quail, respectively, Figure 3b).

### 3.3. Effect of chronic stress during the displacement test

Analyses on the number of errors made before reaching the target cups (either the spatial cup or the displaced cup) showed that animals were more performant to reach the displaced cup, independent of their treatment group. There was also a significant interaction between the test occasion and the target cups (effect of target cup: F_1,35_=100.71, p<0.001; effect of test occasion: F_2,70_=3.22, p=0.046; effect of stress: F_1,35_=0.010, p=0.92; interaction target cup x stress: F_1,35_=0.27, p=0.6; interaction target cup x test occasion: F_2,70_=8.26, p=0.001; interaction target cup x test occasion x stress: F_2,70_=0.87, p=0.42). Post-hoc analyses revealed animals did significantly fewer errors before reaching the displaced cup in all test occasions, compared to the errors before reaching the spatial cup (p=0.001, p<0.001 and p<0.001, for the tests performed after initial training, overtraining, and just after the emotional challenge, respectively, Figure 4a). Between test occasions and within-target cup, post-hoc comparisons revealed that animals made significantly fewer errors before reaching the displaced cup during the tests after overtraining and after the emotional challenge, compared to the test after initial training (p=0.001 and p=0.005, respectively, Figure 4a).

**Figure 4.**
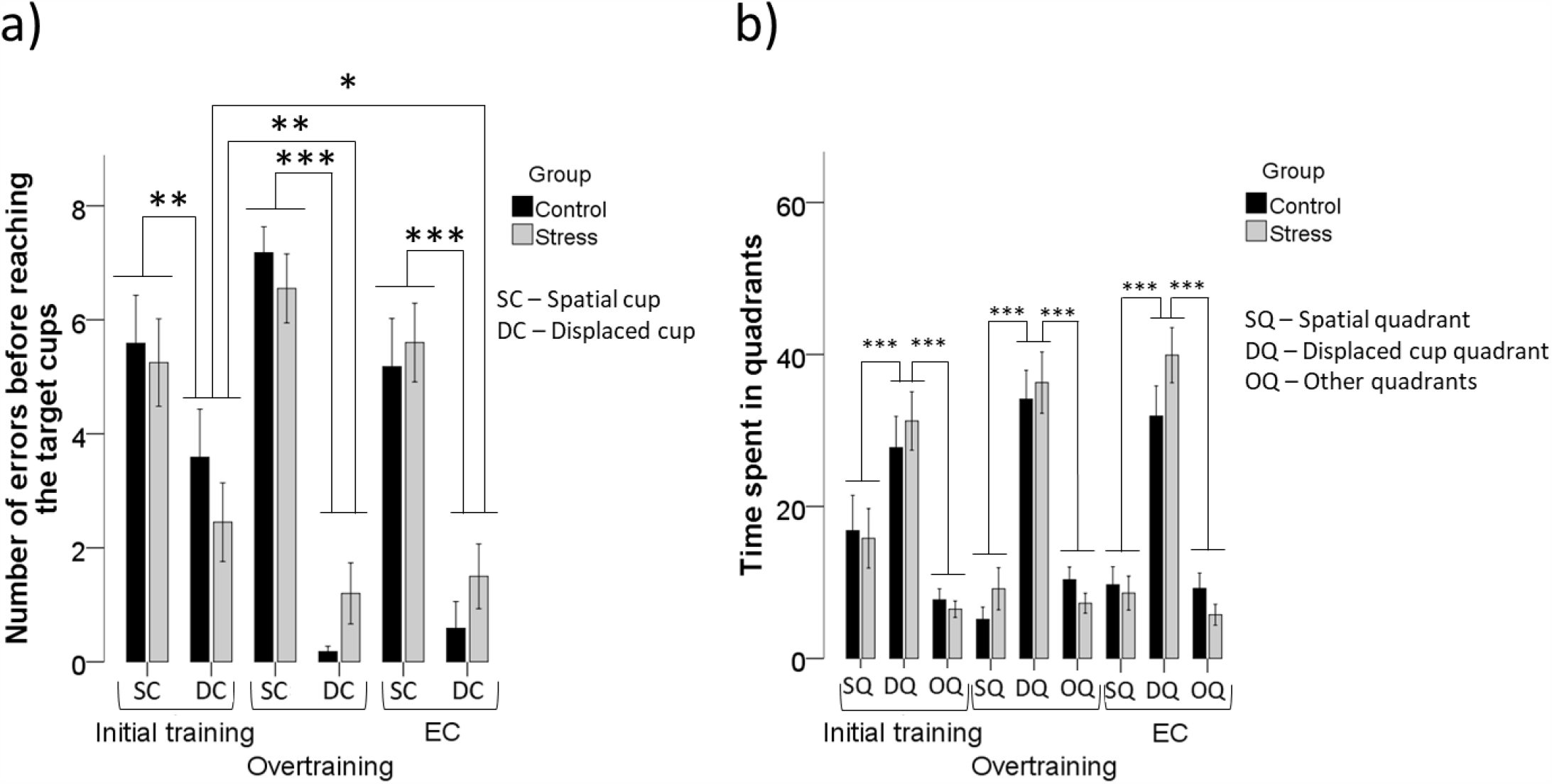
Effects of chronic stress in the displacement tests performed after initial training (5 days), overtraining (15 days), and after an emotional challenge (EC, exposure to an open field). **a)** Number of errors before reaching the target cups (spatial cup or displaced cup), and **b)** Time (in seconds) spent in different quadrants of the arena (either the spatial quadrant, SQ, the displaced cup quadrant, DQ, or the other quadrants, OQ). * p≤0.05; ** p≤0.01; *** p≤0.001, significant difference between the spatial cup and displaced cup or between spatial quadrant, displaced cup quadrant, and other quadrants. Mean ± SEM are given.

Analyses on time spent in different quadrants showed that animals did not spend their time evenly between quadrants. They spent significantly more time than the chance level in the displaced cup quadrant independent of their treatment group. There was also significant a significant interaction between the test occasion and the time spent in different quadrants (effect of quadrants: F_2,70_=67.19, p<0.001; effect of test occasion: F_2,70_=0.82, p=0.44; effect of stress: F_1,35_=3.81, p=0.059; interaction quadrants x stress: F_2,70_=1.09, p=0.34; interaction quadrants x test occasion: F_4,140_=3.25, p=0.014. interaction quadrants x test occasion x stress: F_4,140_=0.42, p=0.78). Post-hoc analyses revealed that the time spent in the displaced cup quadrant was higher for all test occasions compared to those spent on the spatial quadrant and other quadrants (p<0.05, for the tests performed after initial training, overtraining, and just after the emotional challenge, respectively, Figure 4b).

### 3.4. Effect of chronic stress during the colour test

During the colour test, independently of chronic stress, the number of visits made before reaching a cup was different depending on the cup considered (effect of cup: F_2,70_=18.60, p<0.0001; effect of stress: F_1,35_=0.03, p=0.87; interaction cup x stress: F_2,70_=0.06, p=0.95; Figure 5). Post-hoc analyses showed that the number of visits before reaching the spatial black cup was significantly lower than those performed before reaching other black cups or the white cup (p=0.02 and p<0.001, respectively, Figure 5a).

**Figure 5.**
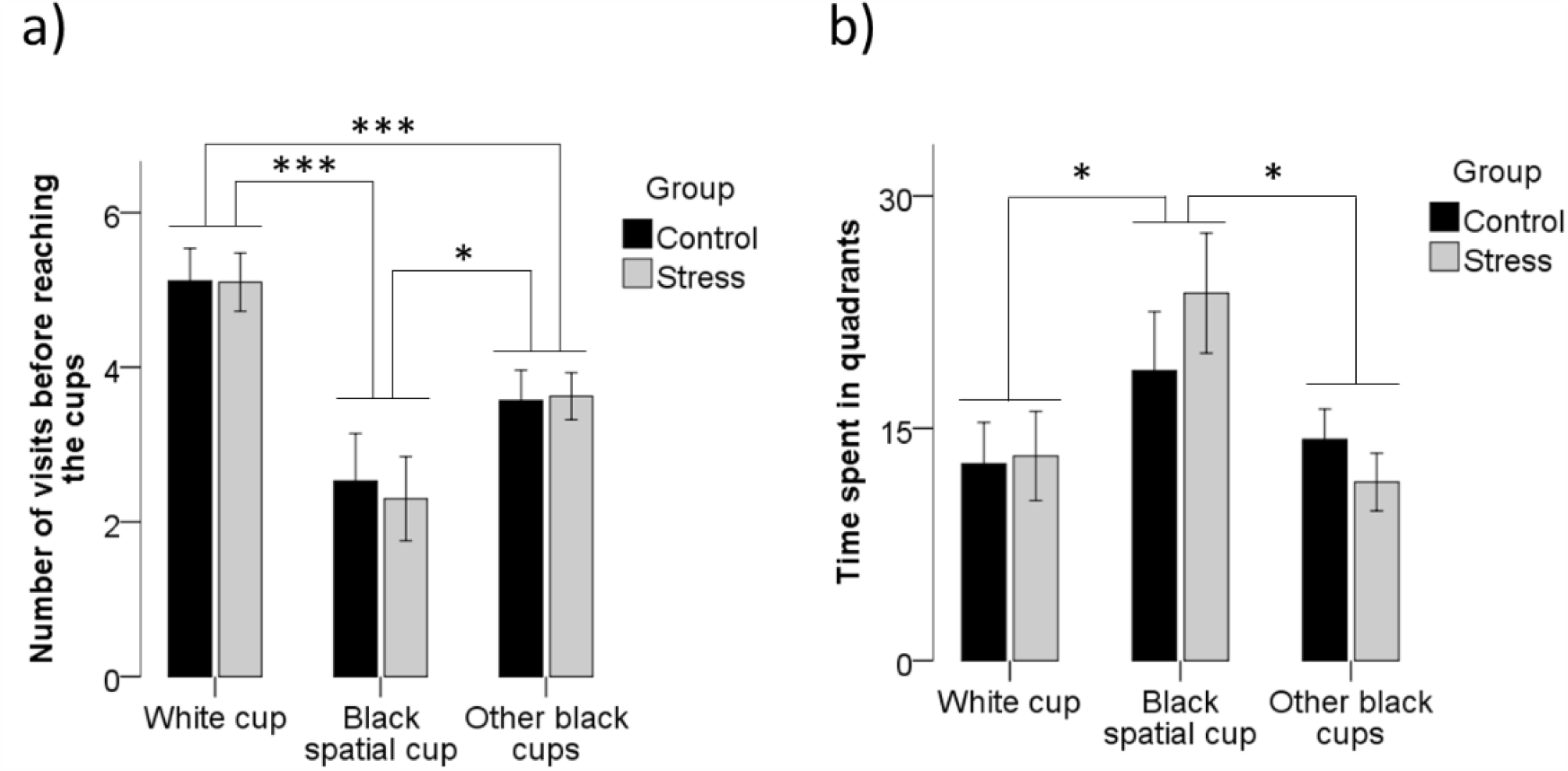
Effects of chronic stress on the colour test. **a)** Number of visits before reaching the white cup, the black spatial cup, or other black cups in chronically stressed and control birds and **b)** Time (in seconds) spent in different quadrants of the arena containing the white cup, the spatial black cup, other black cups in chronically stressed and control birds. * p≤0.05; ** p≤0.01; *** p≤0.001, significant difference between the white cup, black spatial cup, and other black cups. Mean ± SEM are given.

Finally, the time spent in the different quadrants was also significantly dependent on the quadrant considered (effect of quadrants: F_2,70_=3.62, p=0.032). Post-hoc analyses revealed a significantly larger amount of time spent in the black spatial cup quadrant compared to other black cups and white cup quadrants (p=0.01). No differences were found between the treatment groups, nor for the interaction quadrants and treatment groups (effect of stress: F_1,35_=1.02, p=0.31; interaction quadrants x stress: F_2,70_=0.59, p=0.55, Figure 5b).

## 4. DISCUSSION

The main objective of the present study was to address, in birds, whether chronic stress can differentially impact two forms of long-term memory, the declarative spatial and the non-declarative cue-based memories, according to the level of training. To this end, control and chronically stressed Japanese quail were trained in a task that could be solved using spatial and cue-based memory, the dual spatial/cued task, and tested for their performances on each type of memory after 5 and 15 days of training (initial training and overtraining, respectively). Our results showed that the spatial memory of quail was sensitive to the deleterious effect of chronic stress. Performances of chronically stressed birds during the spatial test after initial training were impaired compared to that of control birds. During the test after overtraining, however, these differences were lowered. Interestingly, during the test after an emotional challenge, when birds were exposed to an open field, the negative impact of chronic stress on spatial memory performances was fully reinstated. These findings indicate that the relationships between chronic stress and spatial memory are complex and dynamic. By contrast, cue-based memory was not affected in chronically stressed birds compared to control birds on any test occasion. Thus, this form of memory is relatively resistant to the negative effect of chronic stress. Finally, a test conducted to better qualify birds’ cue-based memory performances, the colour test, provided support that chronically stressed birds were still able to efficiently remember the spatial location of the target cup when this location was signalled by a colour similar to that used during training. This suggests that chronically stressed birds cannot efficiently recall stored spatial information in a context different from the one experienced during training, which highlights that chronic stress reduces the ability to use spatial memory flexibly.

Our results from habituation showed that the number of mealworms eaten increased over days without any significant effect of chronic stress. Moreover, the latency to reach the target cup decreased over days of training in both groups. These findings indicate that both control and chronically stressed birds similarly habituated to the arena and the cups and similarly learned the dual spatial/cued task. They also indicate differences between groups observed during the different tests performed could not be attributed to evident motor or motivational differences and instead reflect stress-related changes in memory performances.

During the spatial test, the number of errors before reaching the spatial location of the target cup was higher in all test occasions (initial training, overtraining, after an emotional challenge) for chronically stressed birds compared to control birds, evidencing the negative impact of chronic stress on spatial memory performances. Although not significant (p=0.07), the interaction between test occasion and stress suggests that the performance of both groups during overtraining tests was similar, compared to initial training and after emotional challenge tests, when their performances differed more. Confirming this tendency, the time spent in different quadrants revealed some specificities of the animals’ performance over the different test occasions. During spatial tests performed after initial training, chronically stressed birds spent their time evenly between the quadrants, while control birds spent significantly more time in the spatial quadrant. The same did not occur after overtraining. Indeed, performances of both control and chronically stressed birds were relatively low during the spatial test performed after overtraining. Contrasting to the spatial tests, chronic stress did not affect cue-based memory performances in all displacement tests performed. In all test occasions, chronically stressed birds and control had the same performance and followed the displaced black cup preferentially. These findings indicate that chronic stress differentially impacts memory depending on the memory system, with the spatial memory being much more sensitive to the adverse effects of chronic stress than the cue-based memory [1,3,6,36].

Our results suggest that chronic stress has a stronger detrimental effect on spatial memory at early stages compared to the late stages of training. They confirm previous studies in birds and mammals reporting chronic stress that can negatively impact this memory system [1,3,6,7,15]. Moreover, the lowering of chronic stress effects on spatial memory performances after overtraining may be explained by a progressive acquisition of a predominant cue-based strategy to solve the spatial test over days of training. In mammals, it is well documented that the involvement of memory systems is a dynamic process that can vary over time of training. In particular, overtraining in mammals favours cue-based memory [17,18,20,21]. In line with this, the number of errors to reach the black target cup during the displacement tests decreased with training, evidencing an improvement of this type of memory over time. These results confirm the idea that the protective effect of an extended period of training on the negative consequences of chronic stress on memory may be due to a progressive shift toward a non-declarative, cue-based memory system.

An alternative explanation for these results could be that extensive exposure to the task was perceived, by the control animals, as stressful on its own, worsening their performance. For rats, while a 4-day training did not impact brain plasticity, a 14-day training schedule drastically reduced hippocampal neurogenesis [23,37]. If that is the case, it is noteworthy to state that the stress from overtraining compared to the CUS procedure was perceived differently by the animals since the negative effect of chronic stress on spatial memory performances were specifically restored after an emotional challenge. Previous studies highlight that acute stress or stress hormone injections experienced just before testing can induce the re-emergence of an extinguished memory [24,25]. These results suggest that the progressive shift toward a cue-based memory system by control quail after an extending period of training is not due to the elimination of the spatial memory, which can re-emerge depending on the more immediate psychological state of the quail. Similarly, studies on mammals have shown that the shift from spatial memory to cue-based memory with increased training is not due to the elimination of the spatial memory [18,21]. Moreover, even after overtraining, the reappearance of chronic stress adverse effects on the spatial memory of chronically stressed quail could indicate a long-lasting sensibility of this memory system.

Finally, our study highlighted that chronic stress affected specific characteristics of spatial memory but did not erase this form of memory. Indeed, in the last test, the colour test, control, and chronically stressed birds made fewer visits before reaching the black cup located in the same spatial location used during training. Chronically stressed birds seemed thus able to remember the location of the target cup when this cup was black, as during training. It suggests that chronic stress specifically impaired the capacity of birds to remember a previously learned location when this location was no more signalled by the cue (black colour). This capacity is crucial to perform the spatial test efficiently. Numerous studies conducted in birds and mammals evidenced that spatial memory critically requires the hippocampus [11,38–42]. In mammals, the hippocampus was shown to be critically involved in forming distinct memories from events with a high level of similarities (pattern separation) and to re-establish previously acquired memory by using incomplete information as recall cues (pattern completion) [43,44]. During spatial tests, the spatial location was no more signalled by the cue (all cups were white). Birds should re-establish the complete spatial-cued information learned during training (one rewarded black cup among seven non-rewarded white cups) from this incomplete information. Under this framework, chronic stress, which negatively affects the hippocampus’s functioning in quail [15], may impair hippocampus-dependent processing of pattern completion.

In conclusion, the present study presents evidence that chronic stress can negatively impact the birds’ declarative spatial memory. It highlights that chronic stress specifically alters the ability to use spatial memory flexibly. Also, we show for the first time in birds that an extended period of training can lower the spatial memory differences between control and chronically stressed individuals, either by a progressive shift to cue-based memory or by the fact that overtraining can be in itself perceived as stressful by control animals, hindering their performance. These hypotheses should be tested further. An emotional challenge, before testing, was capable of reinstating the differences in performance between chronically stressed and control quail showing the long-lasting sensibility of the spatial memory system. Altogether these findings reveal an original and dynamic dialogue between stress and memory systems in birds.

## Supporting information

Raw data

## ACKNOWLEDGEMENTS

This work was supported by the Institut National de Recherche pour l’Agriculture, l’Alimentation et l’Environnement (INRAE). FL and VHBF were supported by a grant from the French Region Centre Val de Loire and from JUNIA ISA, respectively. We are grateful to all the members of the experimental unit PEAT of INRAE (UE PEAT, INRAE, 2018. Experimental Poultry Facility, DOI: 10.15454/1.5572326250887292E12) and particularly to I. Grimaud for providing care to the birds.

## BIBLIOGRAPHY

1. Conrad CD. 2010 A critical review of chronic stress effects on spatial learning and memory. Prog. Neuro-Psychopharmacology Biol. Psychiatry 34, 742–755. (doi:10.1016/j.pnpbp.2009.11.003)

2. Luksys G, Sandi C. 2011 Neural mechanisms and computations underlying stress effects on learning and memory. Curr. Opin. Neurobiol. 21, 502–508. (doi:10.1016/j.conb.2011.03.003)

3. Sandi C. 2013 Stress and cognition. Wiley Interdiscip. Rev. Cogn. Sci. 4, 245–261. (doi:10.1002/wcs.1222)

4. Conboy L, Sandi C. 2010 Stress at Learning Facilitates Memory Formation by Regulating AMPA Receptor Trafficking Through a Glucocorticoid Action. Neuropsychopharmacology 35, 674–685. (doi:10.1038/npp.2009.172)

5. Conrad CD, Magariños AM, LeDoux JE, McEwen BS. 1999 Repeated restraint stress facilitates fear conditioning independently of causing hippocampal CA3 dendritic atrophy. Behav. Neurosci. 113, 902–913. (doi:10.1037/0735-7044.113.5.902)

6. Goodman J, McIntyre CK. 2017 Impaired spatial memory and enhanced habit memory in a rat model of post-traumatic stress disorder. Front. Pharmacol. 8, 1–8. (doi:10.3389/fphar.2017.00663)

7. Moreira PS, Almeida PR, Leite-Almeida H, Sousa N, Costa P. 2016 Impact of chronic stress protocols in learning and memory in rodents: Systematic review and meta-analysis. PLoS One 11, 1–24. (doi:10.1371/journal.pone.0163245)

8. Wright RL, Conrad CD. 2008 Enriched environment prevents chronic stress-induced spatial learning and memory deficits. Behav. Brain Res. 187, 41–47. (doi:10.1016/j.bbr.2007.08.025.Enriched)

9. Packard MG, Goodman J. 2013 Factors that influence the relative use of multiple memory systems. Hippocampus 23, 1044–1052. (doi:10.1002/hipo.22178)

10. O’keefe J, Nadel L. 1978 The hippocampus as a cognitive map. Oxford: Clarendon Press. (doi:10.1016/j.neuron.2015.06.013)

11. Morris RGM, Garrud P, Rawlins JNP, O’Keefe J. 1982 Place navigation impaired in rats with hippocampal lesions. Nature 297, 681–683. (doi:10.1038/297681a0)

12. White NM, McDonald RJ. 2002 Multiple Parallel Memory Systems in the Brain of the Rat. Neurobiol. Learn. Mem. 77, 125–184. (doi:10.1006/nlme.2001.4008)

13. Lormant F, Cornilleau F, Constantin P, Meurisse M, Lansade L, Leterrier C, Lévy F, Calandreau L. 2020 Research Note: Role of the hippocampus in spatial memory in Japanese quail. Poult. Sci. 99, 61–66. (doi:10.3382/ps/pez507)

14. Conrad CD. 2008 Chronic stress-induced hippocampal vulnerability: the glucocorticoid vulnerability hypothesis. Rev. Neurosci. 19, 395–411.

15. Lormant F et al. 2020 Emotionality modulates the impact of chronic stress on memory and neurogenesis in birds. Sci. Rep. 10, 14620. (doi:10.1038/s41598-020-71680-w)

16. Miracle AD, Brace MF, Huyck KD, Singler SA, Wellman CL. 2006 Chronic stress impairs recall of extinction of conditioned fear. Neurobiol. Learn. Mem. 85, 213–218. (doi:10.1016/j.nlm.2005.10.005)

17. Hicks LH. 1964 Effects of Overtraining on Acquisition and Reversal of Place and Response Learning. Psychol. Rep. 15, 459–462. (doi:10.2466/pr0.1964.15.2.459)

18. Packard MG. 2009 Anxiety, cognition, and habit: A multiple memory systems perspective. Brain Res. 1293, 121–128. (doi:10.1016/j.brainres.2009.03.029)

19. Packard MG. 2009 Exhumed from thought: Basal ganglia and response learning in the plus-maze. Behav. Brain Res. 199, 24–31. (doi:10.1016/j.bbr.2008.12.013)

20. Yin HH, Knowlton BJ. 2006 The role of the basal ganglia in habit formation. Nat. Rev. Neurosci. 7, 464–476. (doi:10.1038/nrn1919)

21. Packard MG, McGaugh JL. 1996 Inactivation of hippocampus or caudate nucleus with lidocaine differentially affects expression of place and response learning. Neurobiol. Learn. Mem. 65, 65–72. (doi:10.1006/nlme.1996.0007)

22. Ehninger D, Kempermann G. 2006 Paradoxical effects of learning the Morris water maze on adult hippocampal neurogenesis in mice may be explained by a combination of stress and physical activity. Genes, Brain Behav. 5, 29–39. (doi:10.1111/j.1601-183X.2005.00129.x)

23. Aztiria E, Capodieci G, Arancio L, Leanza G. 2007 Extensive training in a maze task reduces neurogenesis in the adult rat dentate gyrus probably as a result of stress. Neurosci. Lett. 416, 133–137. (doi:10.1016/j.neulet.2007.01.069)

24. Deschaux O, Zheng X, Lavigne J, Nachon O, Cleren C, Moreau JL, Garcia R. 2013 Post-extinction fluoxetine treatment prevents stress-induced reemergence of extinguished fear. Psychopharmacology (Berl). 225, 209–216. (doi:10.1007/s00213-012-2806-x)

25. Izquierdo LA, Barros DM, Medina JH, Izquierdo I. 2002 Stress hormones enhance retrieval of fear conditioning acquired either one day or many months before. Behav. Pharmacol. 13, 203– 213. (doi:10.1097/00008877-200205000-00003)

26. Calandreau L, Bertin A, Boissy A, Arnould C, Constantin P, Desmedt A, Guémené D, Nowak R, Leterrier C. 2011 Effect of one week of stress on emotional reactivity and learning and memory performances in Japanese quail. Behav. Brain Res. 217, 104–110. (doi:10.1016/j.bbr.2010.10.004)

27. Favreau-Peigné A et al. 2016 Unpredictable and repeated negative stimuli increased emotional reactivity in male quail. Appl. Anim. Behav. Sci. 183, 86–94. (doi:10.1016/j.applanim.2016.07.010)

28. Favreau-Peigné A et al. 2014 Emotionality modulates the effect of chronic stress on feeding behaviour in birds. PLoS One 9, 1–12. (doi:10.1371/journal.pone.0087249)

29. Guibert F, Richard-Yris MA, Lumineau S, Kotrschal K, Bertin A, Petton C, Möstl E, Houdelier C. 2011 Unpredictable mild stressors on laying females influence the composition of Japanese quail eggs and offspring’s phenotype. Appl. Anim. Behav. Sci. 132, 51–60. (doi:10.1016/j.applanim.2011.03.012)

30. Laurence A, Lumineau S, Calandreau L, Arnould C, Leterrier C, Boissy A, Houdelier C. 2014 Short-and long-term effects of unpredictable repeated negative stimuli on Japanese quail’s fear of humans. PLoS One 9, 1–8. (doi:10.1371/journal.pone.0093259)

31. Ferreira VHB et al. 2019 Relationship between ranging behavior and spatial memory of free-range chickens. Behav. Processes 166, 103888. (doi:10.1016/j.beproc.2019.103888)

32. Ferreira VHB, Barbarat M, Lormant F, Germain K, Brachet M, Løvlie H, Calandreau L, Guesdon V. 2020 Social motivation and the use of distal, but not local, featural cues are related to ranging behavior in free-range chickens (Gallus gallus domesticus). Anim. Cogn. 23, 769– 780. (doi:10.1007/s10071-020-01389-w)

33. Lormant F, Cornilleau F, Constantin P, Meurisse M, Lansade L, Leterrier C, Lévy F, Calandreau L. 2018 A trait for a high emotionality favors spatial memory to the detriment of cue-based memory in Japanese quail. Behav. Processes 157, 256–262. (doi:10.1016/j.beproc.2018.10.006)

34. Boulay J, Chaillou E, Bertin A, Constantin P, Arnould C, Leterrier C, Calandreau L. 2013 A higher inherent trait for fearfulness is associated with increased anxiety-like behaviours and diazepam sensitivity in Japanese quail. Behav. Brain Res. 237, 124–128. (doi:10.1016/j.bbr.2012.09.026)

35. Parois S, Calandreau L, Kraimi N, Gabriel I, Leterrier C. 2017 The influence of a probiotic supplementation on memory in quail suggests a role of gut microbiota on cognitive abilities in birds. Behav. Brain Res. 331, 47–53. (doi:10.1016/j.bbr.2017.05.022)

36. De Quervain D, Schwabe L, Roozendaal B. 2016 Stress, glucocorticoids and memory: Implications for treating fear-related disorders. Nat. Rev. Neurosci. 18, 7–19. (doi:10.1038/nrn.2016.155)

37. Mohapel P, Mundt-Petersen K, Brundin P, Frielingsdorf H. 2006 Working memory training decreases hippocampal neurogenesis. Neuroscience 142, 609–613. (doi:10.1016/j.neuroscience.2006.07.033)

38. Colombo M, Broadbent NJ, Taylor CSR, Frost N. 2001 The role of the avian hippocampus in orientation in space and time. Brain Res. 919, 292–301. (doi:10.1016/S0006-8993(01)03050-5)

39. Fremouw T, Jackson-Smith P, Kesner RP. 1997 Impaired place learning and unimpaired cue learning in hippocampal-lesioned pigeons. Behav. Neurosci. 111, 963–975. (doi:10.1037/0735-7044.111.5.955)

40. Hampton RR, Shettleworth SJ. 1996 Hippocampal lesions impair memory for location but not color in passerine birds. Behav. Neurosci. 110, 831–835. (doi:10.1037/0735-7044.110.4.831)

41. Kahn MC, Bingman VP. 2009 Avian hippocampal role in space and content memory. Eur. J. Neurosci. 30, 1900–1908. (doi:10.1111/j.1460-9568.2009.06979.x)

42. Squire LR. 1992 Memory and the hippocampus: A synthesis from findings with rats, monkeys, and humans. Psychol. Rev. 99, 582–582. (doi:10.1037/0033-295X.99.3.582)

43. Johnston ST, Shtrahman M, Parylak S, Gonçalves JT, Gage FH. 2016 Paradox of pattern separation and adult neurogenesis: A dual role for new neurons balancing memory resolution and robustness. Neurobiol. Learn. Mem. 129, 60–68. (doi:10.1016/j.nlm.2015.10.013)

44. Neunuebel JP, Knierim JJ. 2014 CA3 retrieves coherent representations from degraded input: Direct evidence for CA3 pattern completion and dentate gyrus pattern separation. Neuron 81, 416–427. (doi:10.1016/j.neuron.2013.11.017)

